# New record of *Dendronephthya hemprichi* (Family: Nephtheidae) from Mediterranean, Israel- an evidence for tropicalization?

**DOI:** 10.1101/2023.07.04.547739

**Authors:** Hagai Nativ, Ori Galili, Ricardo Almuly, Shai Einbinder, Dan Tchernov, Tali Mass

## Abstract

Bioinvasions have the potential to provoke cascade effects that can disrupt natural ecosystems, and cause ecological regime shifts. The Mediterranean Sea is particularly prone to bioinvasions as the changing water conditions, evoked by climate change, are creating advantageous conditions for Lessepsian migrants from the Red Sea. Recently, in May 2023, a new alien species was documented in the Mediterranean Sea - a soft coral of the genus *Dendronephthya*. This discovery was made by divers conducting ‘Long Term Ecological Research’ surveys, along the coast of Israel, at a depth of 42 m. Genetic and morphological testing were utilized to confirm the species identity as *Dendronepthya hemprichi*, an Indo-Pacific coral, common in the Red Sea. According to life history traits of this species such as accelerated attachment to available surfaces and fast growth, we expect it to rapidly expand its distribution and abundance across the Mediterranean.

**Highlights:**

- Potential ‘Tropicalization’ of the Mediterranean Sea
- Increasing water temperatures are an important vector for marine bioinvasion
- A second soft coral species, associated with Lessepsian migration, was identified in the Mediterranean
- The importance of a long-term ecological monitoring program for identifying ecosystem changes

## Introduction

Bioinvasions are some of the most deleterious and pervasive consequences of anthropogenic global change. These invasions can provoke cascade effects that can disrupt natural ecosystems, and cause ecological regime shifts (Conversi et al., 2015). Given suitable environmental conditions and the fragility of an ecosystem, an introduced (alien) species has the potential to become invasive, transforming into a pest within its new habitat and spreading rapidly (Ehrenfeld, 2010). Temperature, in particular, which is highly influenced by climate change, is regarded as a crucial factor that can act as a selective filter, ultimately governing the potential success of alien marine species (Theoharides and Dukes, 2007). Reproductive strategies also play a critical role in the success of sessile benthic invertebrates bioinvasions (Capel et al., 2017). Mixed species assemblages are often generated through sexual reproduction (Hoeksema and Benzoni, 2013), while monospecific aggregations result from asexual reproduction such as budding or fragmentation (Hoeksema, 2004; Hoeksema et al., 2019; Hoeksema and Gittenberger, 2010). Reproduction strategies are pivotal to enabling invasive species to out-compete native species through overgrowth, smothering, or competitive exclusion. For instance, *Tubastraea* spp. employ minor but continuous sexual reproduction and clonal reproduction, enabling them to dominate large, rocky surfaces and displace native corals and zoantharians (Creed, 2006; Luz and Kitahara, 2017; Miranda et al., 2016).

Marine traffic and man-made canals are some of the main pathways for the introduction of marine invasive species (Carlton, 1985; Gollasch et al., 2006; Hewitt et al., 2009). One example is the Suez Canal, which is the primary pathway for the introduction of over half of the non-indigenous species in the Mediterranean Sea (Galil et al., 2018; Por, 2012).

The Mediterranean Sea is an ultra-oligotrophic (Krom et al., 2004) semi-enclosed temperate sea, which exhibits a high salinity of 39 ppt (Soukissian et al., 2017), a wide annual temperature range (15°C - 30°C) (Saraçoğlu et al., 2021; Shaltout and Omstedt, 2014). The Mediterranean Sea is unique in its rapidly changing ecosystems, affected by both climate change and the introduction of invasive species. More specifically, the eastern Mediterranean region is experiencing rapid warming due to climate change, and in recent decades, water temperatures have been rising at a rate of 0.35 ± 0.27 °C decade^-1^ (El-Geziry, 2021; Sisma-Ventura et al., 2014). These trends also result in an increase of the minimum winter temperatures in coastal waters, with minimum temperatures shifting from 16°C to 18°C since the 1990s (Ozer et al., 2017). Collectively, these temperature shifts may provide favorable conditions for the invasion of warm-water species (Ehrenfeld, 2010).

Across the Mediterranean Sea, over 500 alien species originating from the Red Sea, have been documented to date (Galil, 2007; Galil et al., 2014; Golani, 1998; Goren and Galil, 2005; Spanier and Galil, 1991), a phenomenon known as ‘Lessepsian migration’(Por, 2012). These alien species (fish, invertebrates and algae) have arrived via the Suez Canal, which has undergone several expansion projects since it was first dredged in 1869, thereby removing the depth and salinity barriers which once hindered the crossing for various species (Castellanos-Galindo et al., 2020). Israel’s coast has often been documented as the “first stop” for alien species that later establish stable populations across the entire Mediterranean Sea (Hoffman et al., 2014; Karachle et al., 2004; Kletou et al., 2016). This is due to Israel’s proximity to the Suez Canal, located just to the south, coupled with the prevailing south to north currents that run along Israel’s coast as part of the larger counterclockwise circulation of the Mediterranean Sea (Hamad et al., 2006).

Although various alien species from diverse phyla have been recorded in the Mediterranean, the only documented alien soft coral (Cnidaria: Octocorallia) to date is *Melithaea erythraea* (Ehrenberg, 1834). This soft coral is native to the Red Sea and has arrived in the Mediterranean through Lessepsian migration. *M. erythraea* was first documented in the Hadera power plant (Israel) in 1999, and remained confined within that facility until 2015 when additional colonies were observed on the surrounding rocky reefs (Fine et al., 2005; Grossowicz et al., 2020).

Soft corals are found world-wide, in a wide range of depths and water temperatures (Fabricius and Alderslade, 2001), and are the second most abundant sessile organism in many coral reefs (Benayahu and Loya, 1981). Several species of soft corals are indigenous to the Mediterranean, mainly from the families Pennatuloidea, Alcyoniidae, and Hormathiidae (Barış Özalp and Suat Ateş, 2015; Vafidis et al., 1994). The genus *Dendronephthya* (Cnidaria: Octocorallia: Alcyonaea: Nephtheidae), is a soft coral found in the tropical waters of the Indo-Pacific Ocean. This genus has been documented at a wide range of depths, and is primarily in habitats with strong currents (Fabricius and Alderslade, 2001; Fabricius et al., 1995b; Grossowicz and Benayahu, 2012) such as vertical artificial structures or steep reefs (Perkol-Finkel and Benayahu, 2004). *Dendronephthya* is an azooxanthellate species (Fabricius et al., 1995a), characterized by a wide range of bright colors, eight pinnate tentacles on each polyp, a branching divaricate structure supported by a hydrostatic skeleton, and internal calcareous skeletal elements called sclerites (Dahan and Benayahu, 1997a, 1997b; Pupier et al., 2018). It is a passive suspension feeder, depending on ambient currents for the supply of food particles, mainly phytoplankton (Fabricius et al., 1995a).

*Dendronephthya* tends to rapidly populate available substrate, often artificial surfaces, in as little time as two days (Dahan and Benayahu, 1997b; Fabricius et al., 1995a, 1995b; Perkol-Finkel and Benayahu, 2007, 2004). *Dendronephthya* is fast growing, and (Perkol-Finkel and Benayahu, 2005) report that the number of *Dendronephthya* sp. colonies can increase four-fold in one year following initial recruitment to new habitat. Attachment to artificial structures is advantageous as it enables exposure to high currents, which in turn, enhances the coral’s growth (Perkol-Finkel and Benayahu, 2004). In addition to sexual reproduction, *Dendronephthya* can also reproduce via clonal propagation (Dahan and Benayahu, 1997a, 1997b) where autotomized fragments, which are negatively buoyant, settle on the outer face of a horizontal substratum (Perkol-Finkel and Benayahu, 2007). Due to their fast growth and reproductive strategies, *Dendronephthya* often become the most abundant sessile organisms on these structures, (Perkol-Finkel and Benayahu, 2007, 2005).

Along the Mediterranean coast of Israel, underwater surveys are conducted bi-annually at nine locations (Fig. 1), as part of a Long Term Ecological Research (LTER) program, by the Morris Kahn Marine Research Station (MKMRS). (MKMRS LTER; established in 2014— https://med-lter.haifa.ac.il/index.php/en/data-base). In May 2023, MKMRS LTER researchers observed and documented the first record of *Dendronephthya* sp. in the Mediterranean Sea, during a routine survey.

**Figure 1.**
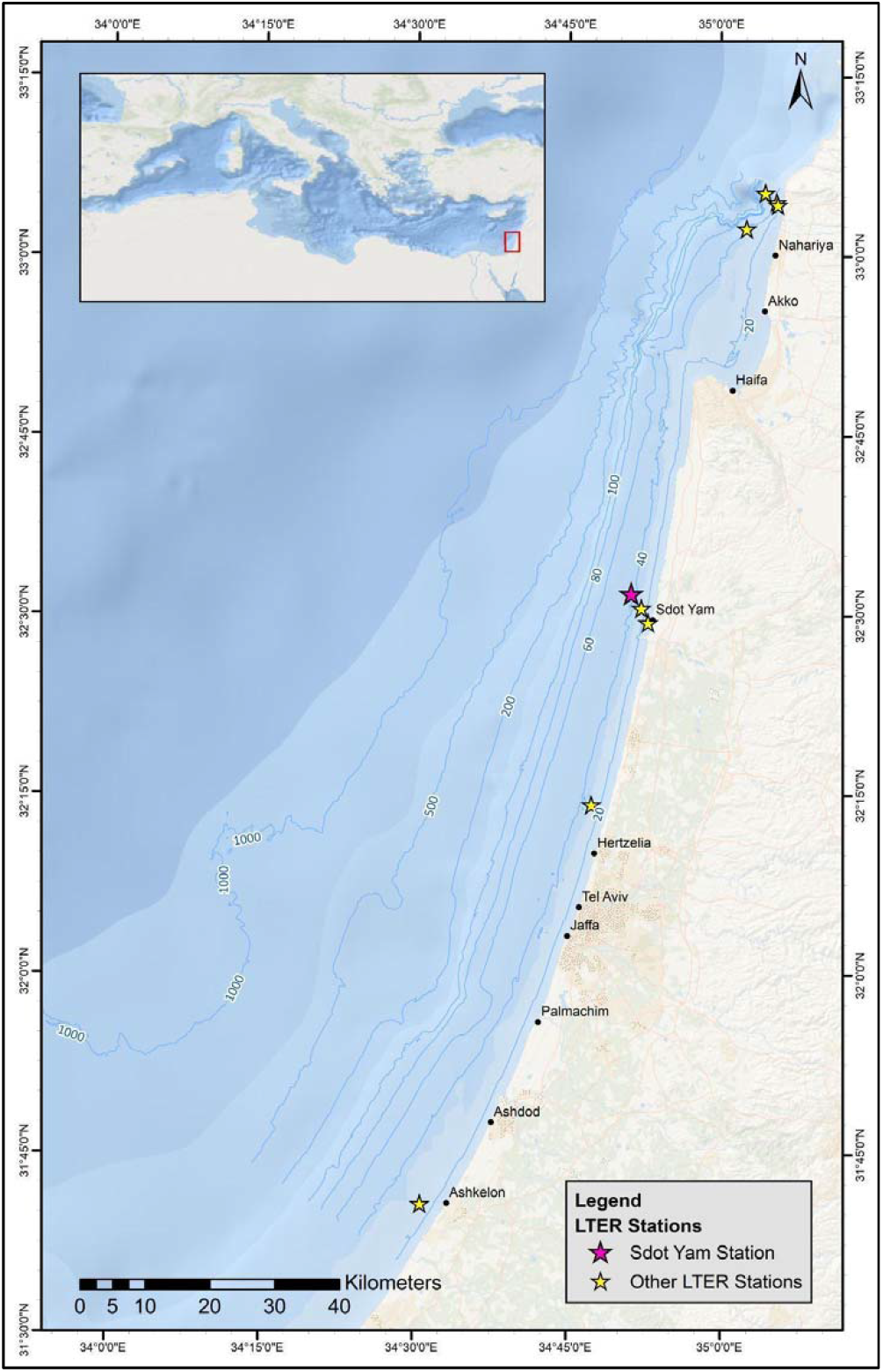
Map of the Israeli Mediterranean coast, and the nine MKMRS LTER monitoring sites.

In this study, we describe *Dendronephthya* sp. with an aim to understand the expansion potential of this species at the Mediterranean Sea.

## Material & Methods

### Study Site and Sample Collection

On May 18, 2023, MKMRS LTER researchers encountered several colonies of *Dendronephthya* sp. along the rocky reef monitoring site (32.54°N, 034.85°E) located near Sdot Yam, Israel, at a depth of 42 m (Fig. 1). This rocky reef site is exposed to open sea currents. At the time of the survey, water temperature was 19°C. Specimens from three colonies were collected from the natural rocky reef in the Mediterranean. In order to compare these specimens with native populations in the Red Sea, seven colonies were later collected, under a special permit from the Israel Nature and Parks Authority, in the Red Sea, adjacent to the Inter-University Institute (29.30°N, 34.54°E), in Eilat, Israel. The seven specimens from the Red Sea were collected from both artificial structures and natural reefs. Four colonies were collected from a depth of 36 m, two of which were associated with artificial structure and two of which with the natural reef. An additional three colonies were collected at a depth of 12 m, and these were associated with artificial structures. Samples from each colony were sectioned into fragments 1 cm^2^ in size, and were flash frozen in liquid nitrogen prior to DNA extraction. The remainder of the specimens collected were utilized for further morphological examination.

### DNA Extraction

Genomic DNA was extracted using a Proteinase-K protocol (Geraghtly et al 2013). For species identification, two genetic markers were PCR-amplified. The mitochondrial marker ribosomal 16S gene was amplified using primers DN1-F (5’-AGGCTACTTAAGTATAGGGG-3’) and DN1-R (5’-AACTCTCCGACAATATTACGC-3’), with PCR conditions described in (Williamson et al., 2022). The second marker was the oxidase subunit II and I (cox1 and 2) which were generated based on the available sequences of *Dendronephthya hemprichi* (native to the Red Sea) in NCBI at the time (GU355996.1), DhCox12F AGAGTGTTCTCACCTACTTTAG, DhCox12R GTTTAGCAGAAAATGTGGGTAT. All PCRs were performed using Kodaq 2X PCR MasterMix (ABM, Richmond, BC Canada) following the manufacturer’s protocol. DNA yield and PCR products were analyzed by electrophoresis on a 1.0% agarose TBE (90 mM TRIS-borate and 2 mM EDTA) gel run at 110 V. PCR products were Sanger sequenced in both directions using the amplification primers on ABI 3730xl DNA Analyzer at HyLabs (Israel). Sequences from each gene were (separately) aligned to complete mitochondrial genomes of the several species of *Dendronephthya* and Octocorallia available in NCBI’s GenBank (Table 1) using MUSCLE algorithm in MEGA11 (Kumar et al., 2018). Alignments were trimmed to retain shared regions among all sequences. The trimmed alignment of the ribosomal 16S gene and cox1 and 2 gene included 547 bases and 830 bases, respectively. For species *D. hemprich*i a reference sequence was found in public databases only for the cox1 gene (NCBI accession number GU355996.1) but no reference for rRNA was available. Then, the ribosomal 16S gene and cox1 gene alignments were concatenated and used for calculation of a maximum likelihood tree, with the PhyML 3.0 algorithm (Guindon et al., 2010), the web application (http://www.atgc-montpellier.fr/phyml/). Standard bootstrap analysis was performed with 1000 repeats.

**Table 1:**
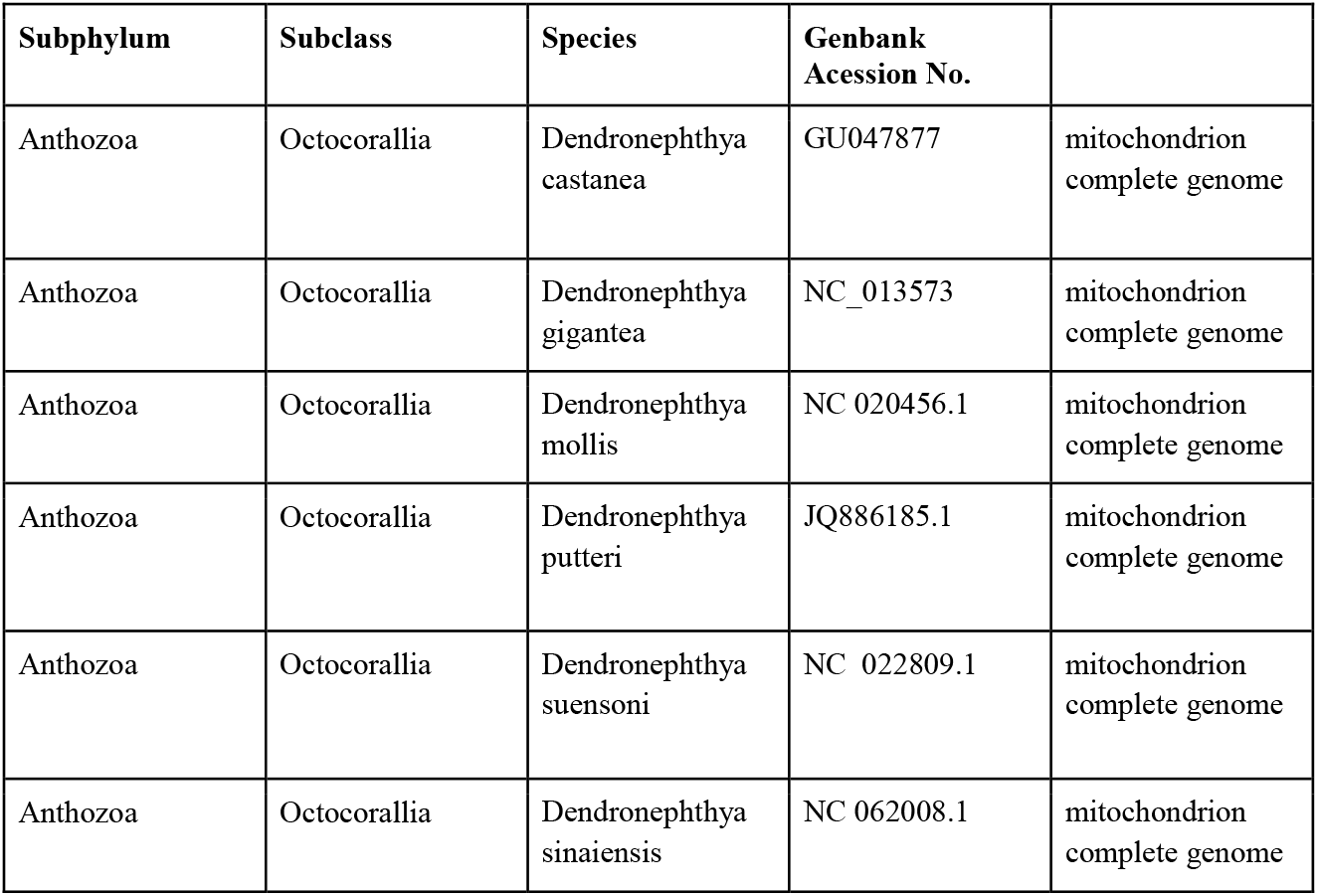
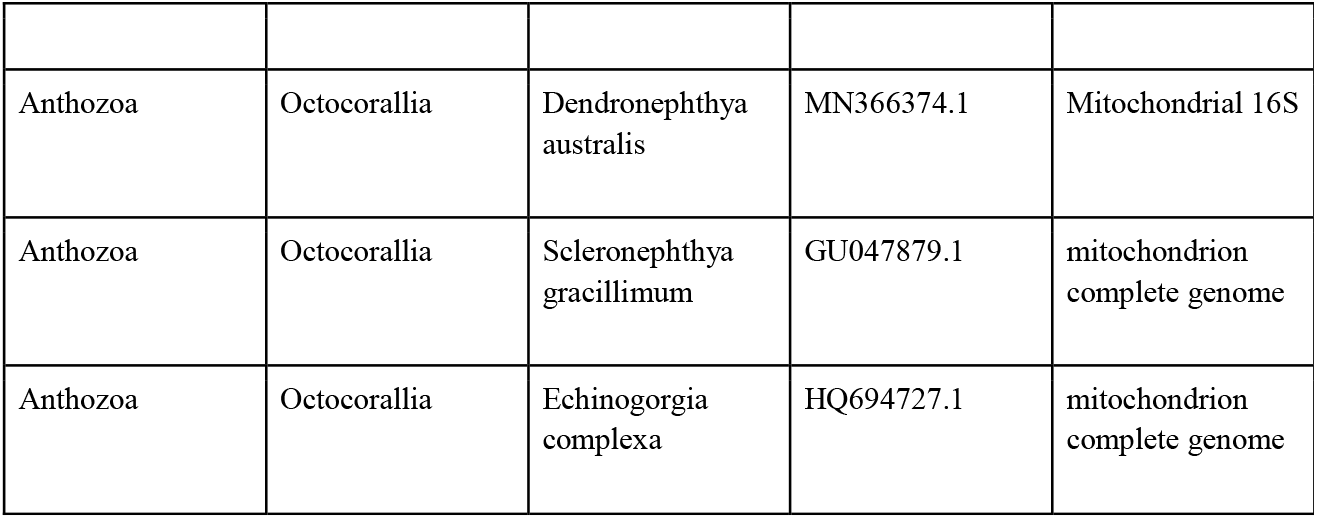
Specimens used for phylogenetic analysis in this study

| Subphylum | Subclass | Species | Genbank<br>Accession No. |  |
| --- | --- | --- | --- | --- |
| Anthozoa | Octocorallia | Dendronephthya<br>castanea | GU047877 | mitochondrion<br>complete genome |
| Anthozoa | Octocorallia | Dendronephthya<br>gigantea | NC_013573 | mitochondrion<br>complete genome |
| Anthozoa | Octocorallia | Dendronephthya<br>mollis | NC 020456.1 | mitochondrion<br>complete genome |
| Anthozoa | Octocorallia | Dendronephthya<br>putteri | JQ886185.1 | mitochondrion<br>complete genome |
| Anthozoa | Octocorallia | Dendronephthya<br>suensoni | NC 022809.1 | mitochondrion<br>complete genome |
| Anthozoa | Octocorallia | Dendronephthya<br>sinaiensis | NC 062008.1 | mitochondrion<br>complete genome |
| Anthozoa | Octocorallia | *Dendronephthya australis* | MN366374.1 | Mitochondrial 16S |
| Anthozoa | Octocorallia | *Scleronephthya gracillimum* | GU047879.1 | mitochondrion complete genome |
| Anthozoa | Octocorallia | *Echinogorgia complexa* | HQ694727.1 | mitochondrion complete genome |

### Sclerite Morphological Analysis

Sclerites were isolated from segments of coral tissue, 1-2 cm in length, using 3% sodium hypochlorite. Once tissue was no longer visible, remaining sclerites were rinsed three times with DDW and stored in 100% EtOH. Sclerites were placed on silica wafer, mounted on SEM plugs and vacuum coated with 5 nm Au/Pb (80:20%) prior to examination under a ZEISS SigmaTM SEM (Germany), by using a SE2 detector (1-2 kV, WD = 6-7 mm). The length and width of the different types of sclerites found in the polyps were measured using Fiji software (Schindelin et al. 2012) (77 sclerites).

## Results

In total, fifteen colonies were observed in a 10 m^2^ area of rocky reef, with colony size ranging from 5 to 50 cm (Fig. 2), suggesting the utilization of a propagation strategy. This surveyed area is one of the nine sites routinely surveyed every year since 2014, in both the spring and fall. Routine surveys include both photo quadrats for assessments of percent cover of algae and invertebrates, as well as fish counts. At the remaining eight rocky reef monitoring stations, during the spring 2023 survey, no evidence of additional colonies of *Dendronephthya* sp. was found. Furthermore, no soft corals were observed during any of the previous survey years (https://med-lter.haifa.ac.il/index.php/en/data-base).

**Figure 2.**
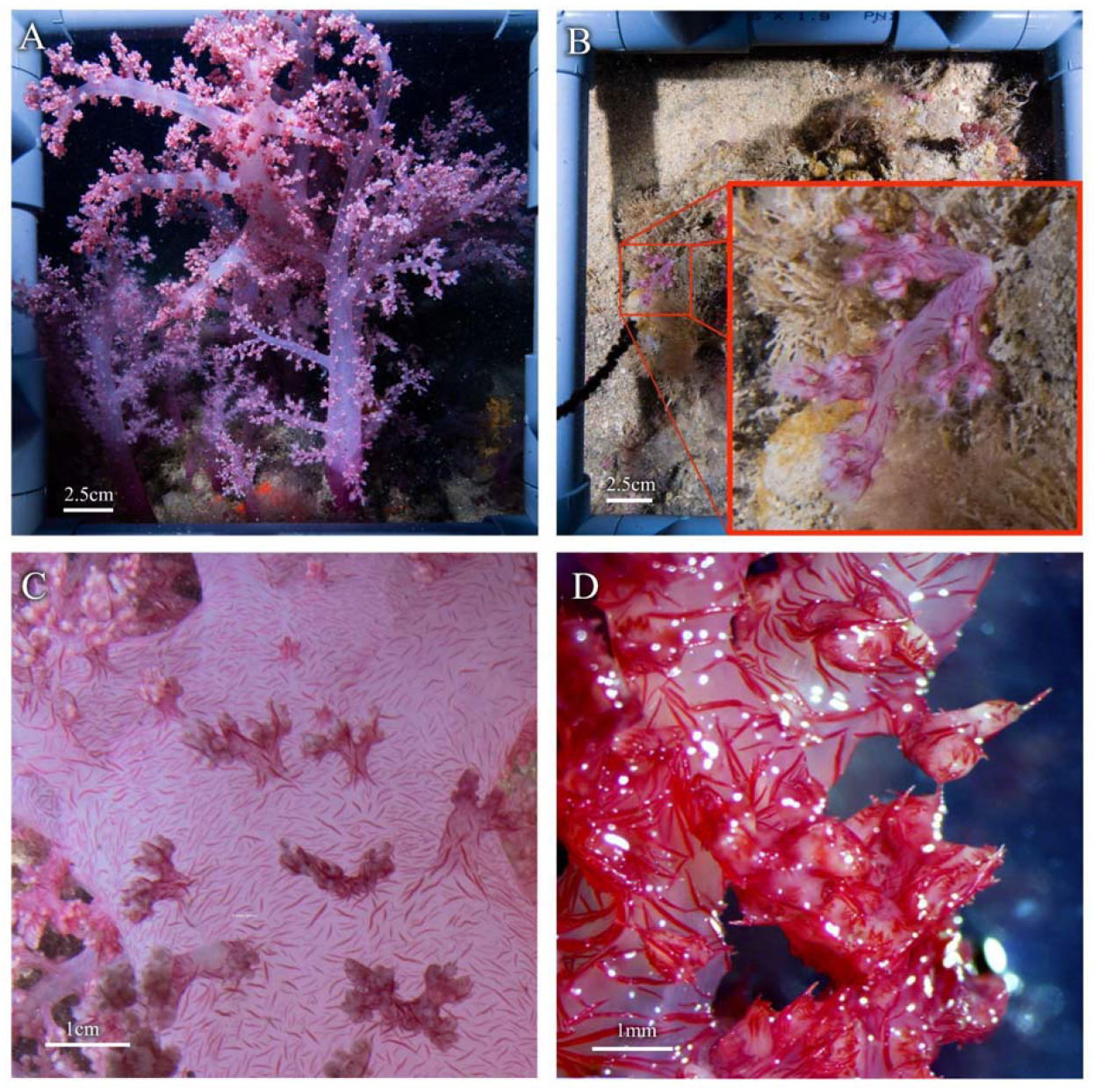
*Dendronephthya hemprichi* colonies observed at the rocky reef at 42 m near Sdot-Yam, Israel. (A-B) Size variation of the colonies. The photographed colonies vary from (A) ∼50 cm to (B) ∼ 5 cm from base to top. (C) Red spindle-shaped sclerites arrangement on the stalk surface. (D) Supporting bundle and loose spindle sclerites arranged around the polyp head.

The observed colonies displayed distinctive traits associated with the genus *Dendronephthya*. These include vivid red pigmentation, as well as their intricate branched divaricate structure. Furthermore, the polyps exhibited a notable arrangement of spindle shaped sclerites in their armature, both within the polyp itself and on the surface of the stalk (Fabricius and Alderslade, 2001). Notably, the polyps displayed a conspicuous supporting bundle of large red spindle-shaped sclerites (Fig 2C). Lastly, similar thick spindle-shaped sclerites were observed on the surface of the stalk (Fig. 2D).

Therefore, to confirm the species identity of the soft corals observed near Sdot-Yam, a phylogenetic analysis using mitochondrial small subunit ribosomal RNA and cox1 and 2 gene sequences was performed. The analysis revealed that all studied colonies belong to the genus *Dendronephthya* (Fig. 3A), and that the colonies collected near Sdot-Yam had high similarity to the colonies collected at the Red Sea.

**Figure 3.**
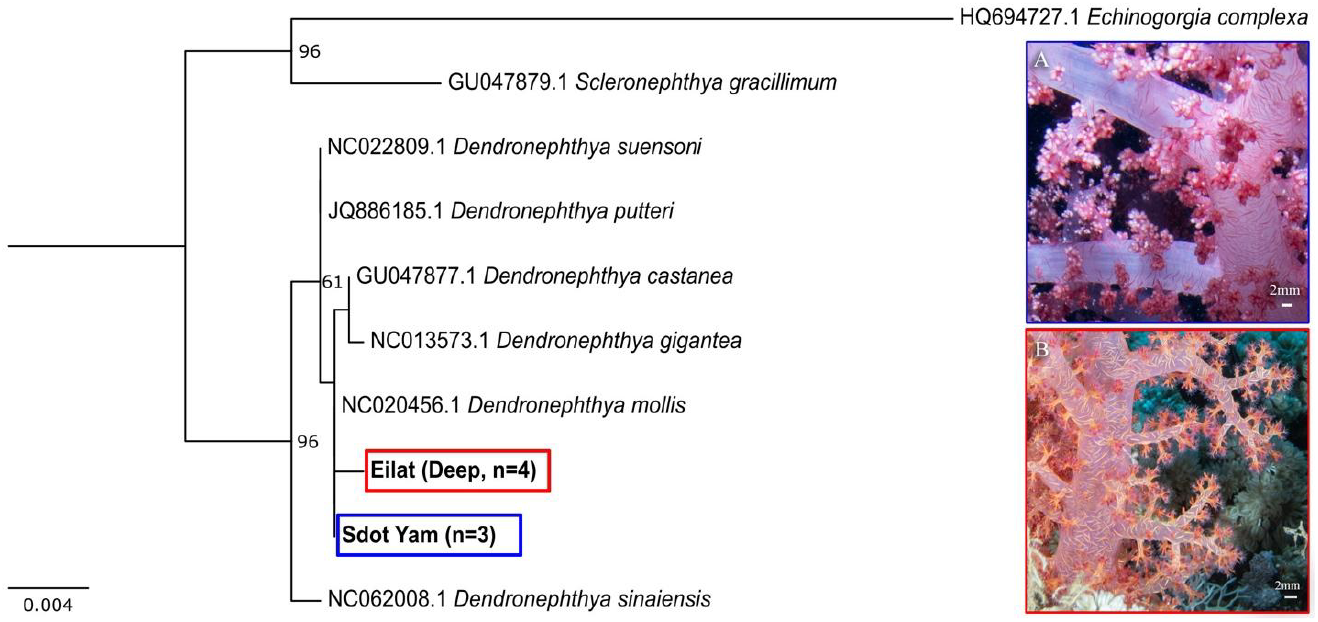
Species identity verification of *Dendronephthya hemprichi*, from colonies collected, using both genomic markers (ribosomal 16S and cox1 gene). A maximum likelihood tree based on sequences of mitochondrial small subunit 16S gene and cox1 gene. Tree was calculated using the PhyML 3.0 algorithm. Numbers at the nodes indicate bootstrap values (percent of 1,000 repeats). Scale bar represents number of substitutions per site.

The two most abundant species at the Red Sea are *Dendronephthya sinaiensis* and *D. hemprichi*, but complete mitochondrial genome sequence is only available for *D. sinaiensis*. To compensate, we examined sequence similarity between *D. hemprichi* cox1 gene, that is publically available for *D. hemprichi* (NCBI accession number GU355996.1) and cox1 gene sequence in our Sdot Yam and Red Sea specimens. *D. hemprichi* cox1 gene sequence showed 100% base pair identity between cox1 sequence of the Sdot-Yam specimens.

Microscopic examination of the sclerite morphology by SEM (Fig. 4) further verified that all samples belong to the species *D. hemprichi*. Specifically, all colonies contain several millimeter long spindle-shaped sclerites, which are typical to the supporting bundle (Fig. 4A-B). Additional sclerites were observed, including shorter spindle-shaped sclerites, up to 500µm, typically found on the polyp head (Fig. 4C-D), and small flattened, irregular shaped sclerites typical to the stalk section of the coral (less than 100 µm) (Fig 4E-F) (Fabricius and Alderslade, 2001; Grossowicz and Benayahu, 2012). The average length-to-width ratio of the spindle-shaped sclerites is 12.68 ± 5.07 (n = 76 sclerites) which is within *D. hemprichi reported* range ration (Grossowicz and Benayahu, 2012).

**Figure 4.**
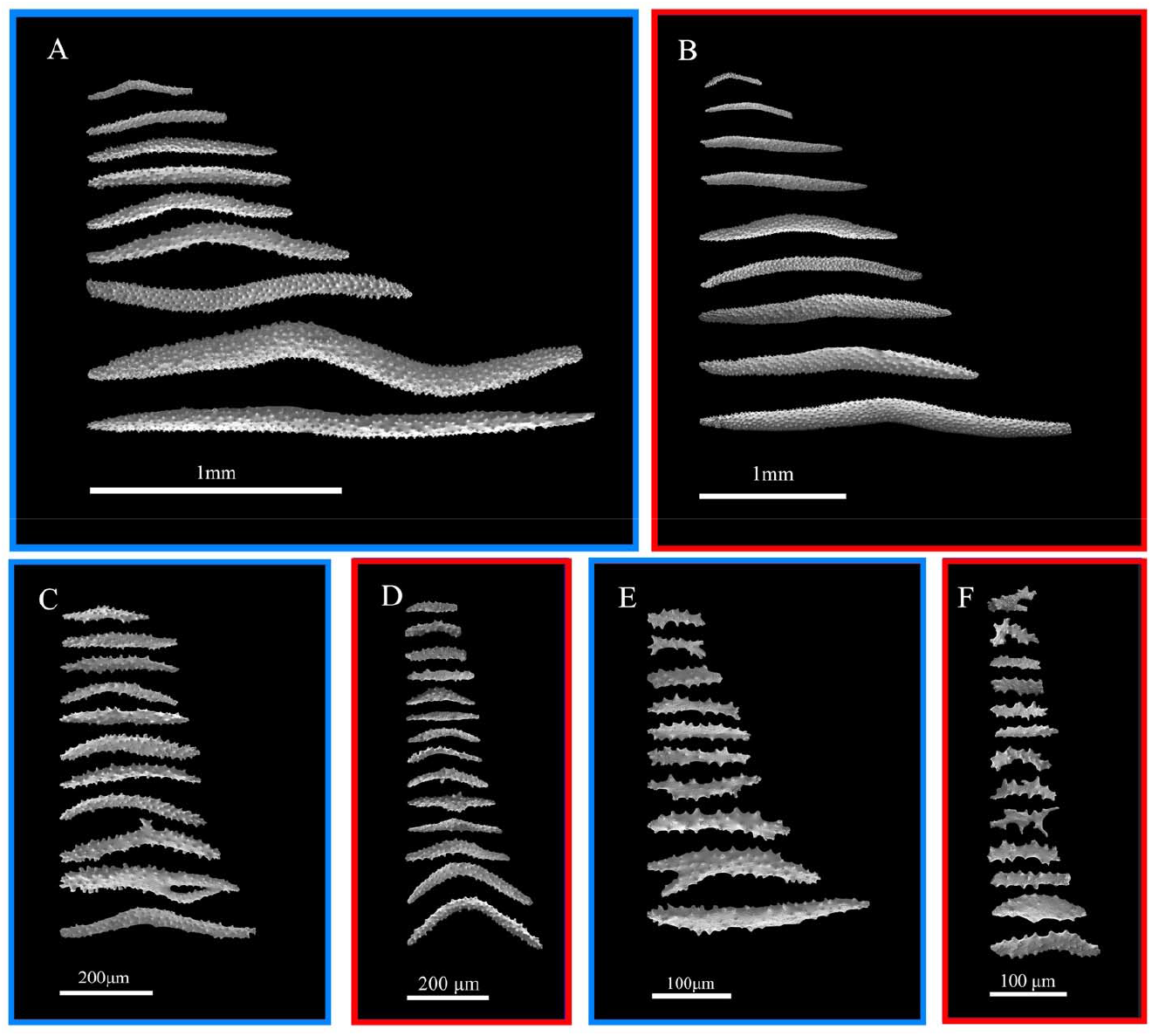
Scanning Electron Micrograph images of spindle-shaped sclerites from the Sdot Yam (blue outline) and Red Sea (red outline) colonies. (A-B) Spindles from the supporting bundles, from colonies collected from the Sdot Yam and Red Sea colonies, respectively. (C-D) Spindles from the polyp heads, from colonies collected from the Sdot Yam and Red Sea colonies, respectively. (E-F) Irregular shaped sclerites from the stalks, from the Sdot Yam and Red Sea colonies, respectively.

## Discussion

Molecular and morphological analysis suggest that the soft coral populations documented in this research belong to the species *D. hemprichi*. To the best of our knowledge, this is the first evidence of occurrence of the Indo-Pacific *D. hemprichi* within the Mediterranean Sea. The genus *Dendronephthya* is common in the northern Red Sea and the Indo-pacific (Fabricius and Alderslade, 2001). The two most common species at the northern Red Sea are *D*.*hemprichi* and *D. sinaiensis. D. hemprichi* is found in a wide range of depths, while *D. sinaiensis* was found to prefer water depths greater than 18 m, with low light intensity (Grossowicz and Benayahu, 2015). The morphological differences which were observed in the polyps of the two species, indicative of differences in feeding niches, mainly related to prey size (Grossowicz and Benayahu, 2012).

Although the exact method of transport can not be determined (asexual autotomized fragments, ship ballast or other), *D. hemprichi* has likely arrived from the Red Sea, via the Suez Canal, as part of the greater and well-documented Lessepsian Migration. Although in its natural habitat, in the Red Sea, *D. hemprichi* is mostly found on artificial vertical structures exposed to high flow regimes (Perkol-Finkel and Benayahu, 2004), the *D. hemprichi* colonies observed in this study, were located at a natural rocky reef site. This interesting observation may indicate that this site experiences high exposure to currents.

To date, only one instance of Lessepsian migration involving an alien soft coral species has been observed in the Mediterranean. The gorgonian species *M. Erythraea*, was first reported by Fine et al. (Fine et al., 2005) in 1999. Despite its reproductive strategy, which confers upon it a substantial capability for swift establishment, the observed restricted expansion of its distribution in the region (Grossowicz et al., 2020), raises the possibility that this species lacking competitive characteristics, or exhibiting limited adaptability to the new ecological conditions encountered in its new habitat.

In contrast, we anticipate a rapid increase in the distribution of *D. hemprichi* throughout the Mediterranean Sea. Previous studies consistently demonstrate its swift colonization of available substrate, capable of multiplying colony numbers four-fold within a single year (Perkol-Finkel and Benayahu, 2007, 2005, 2004). Furthermore, in specific environments, it frequently emerges as the dominant sessile organism, indicating its competitive prowess (Fabricius et al., 1995b). Notably, the clonal propagation model (Dahan and Benayahu, 1997b), characteristic of Dendronephthya species, finds support in the observed diversity of colony sizes within localized areas of the Mediterranean rocky reef. Thus, the future trajectory of *D. hemprichi* in the Mediterranean holds great potential for unraveling its rapid expansion dynamics and ecological significance within the marine ecosystem.

Marine ecosystems are currently facing combined effects from climate change and local human stressors, which have the potential to induce profound shifts at the levels of species, trophic dynamics, habitats, and entire ecosystems. The precise outcome of these interactions varies depending on the specific nature of the interaction itself (Gissi et al., 2021). Lessepsian migration serves as an exemplary illustration of such interplay. The construction of the Suez Canal and subsequent intensified shipping activities have facilitated the arrival of alien species, while climate change has concurrently reshaped the aquatic environment, rendering it conducive for the establishment of viable populations. In fact, some researchers have gone so far as to suggest that the combined effects of climate change and localized human stressors could potentially drive certain local species towards functional extinction (Edelist et al., 2013).

In some studies, the processes observed in the Mediterranean Sea are referred to as ‘Tropicalization’ (Bianchi, 2007; Bianchi and Morri, 2003). This overarching concept encapsulates the overall transition of the region from a temperate ecosystem towards one with tropical characteristics. Such a transformation holds a variety of implications for the biota inhabiting these waters. On one hand, it may offer a competitive advantage to thermophilic species, enabling their proliferation. On the other hand, species with lower thermal tolerance may be compelled to seek refuge in deeper, cooler waters (Martinez et al., 2021) or find themselves operating at the boundaries of their physiological limits. Additionally, there have been reports of poleward range expansions in several cnidarian species (Canning-Clode and Carlton, 2017; Vergés et al., 2014). These findings collectively underscore the complex dynamics at play in response to the combined impacts of climate change and local human stressors within marine ecosystems.

## Conclusion

Once an alien species has migrated to a new environment, its long-term survival and successful establishment as a population rely not only on favorable environmental conditions (Rilov et al., 2009), but also on various additional factors encompassing phenotypic plasticity, competition dynamics, predatorprey interactions, and reproductive strategies (Hewitt et al., 2009). In the case of *D. hemprichi*, this particular species demonstrates several advantageous traits that will likely contribute to its successful establishment in the Mediterranean region, as species originating in the highly biodiverse Red Sea are typically more competitive than species native to the Mediterranean Sea. Furthermore, this soft coral faces minimal predation pressure, and its main mode of propagation involves asexual reproduction (Dahan and Benayahu, 1997a, 1997b), enabling expeditious expansion and growth.

Amidst the doom’s day predictions of a collapsing ecosystem, bio-invasions in the Mediterranean Sea can also be observed through rose-colored glasses. As the Mediterranean Sea is a remnant of the Tethys Ocean (Hewitt et al., 2009), from a historical perspective, the changes observed can also be viewed as a return to the sea’s tropical origins (Por, 2009).

## Acknowledgments

We thank Lalzar from the Bioinformatics Core Unit, the University of Haifa for fruitful discussions and suggested improvements. This research was supported by the Ministry of Innovation, Science & Technology, Israel.

